# Basal activation of ERK1/2 blunts the antimicrobial activity of neutrophils from aged hosts against antibody opsonized *Streptococcus pneumoniae*

**DOI:** 10.1101/2025.05.27.656427

**Authors:** Shaunna R. Simmons, Annabel Rivera, Elsa N. Bou Ghanem

## Abstract

The decline in neutrophil antimicrobial function renders vaccinated aged hosts less protected against *Streptococcus pneumoniae* infection. In vaccinated hosts, activation of neutrophils via phagocytic receptors, like complement and Fcγ receptors, mediates bacterial uptake and killing. However, the mechanisms behind changes in signaling downstream of these receptors with age is not known. Mitogen-activated protein kinases (MAPK) are signaling cascades activated in response to cell surface receptors. Using bone marrow neutrophils isolated from young and aged mice, and MAPK phosphorylation array, we found differences in MAPK activation in an opsonin dependent manner (complement vs antibody). Neutrophils from aged mice failed to increase MAPK phosphorylation upon infection with pneumococci opsonized with heat-inactivated (HI) sera from hosts immunized with the pneumococcal conjugate vaccine (PCV), indicating an age-related defect in FcγR signaling. Neutrophils from aged mice had higher basal levels of MAPK phosphorylation when compared to young controls, including a 15-fold increase in ERK1/2 phosphorylation. Inhibition of ERK1/2 signaling blunted pneumococcal killing by PMNs from young mice, but improved killing by neutrophils from aged mice, only when the bacteria were opsonized with HI immune but not naïve sera. In young human participants, *in vitro* inhibition of ERK1/2 in neutrophils resulted in decreased pneumococcal killing, but only following PCV vaccination, suggesting that the effects of ERK1/2 inhibition are clinically relevant in vaccinated hosts. This study identifies age-related changes in ERK1/2 activation and demonstrates that balanced activation of this pathway is crucial for neutrophil antimicrobial activity against antibody opsonized bacteria following vaccination in both mice and humans.

**Summary sentence:** Imbalanced ERK1/2 activation impairs neutrophil activity in aged hosts.

**LEGENDS:** 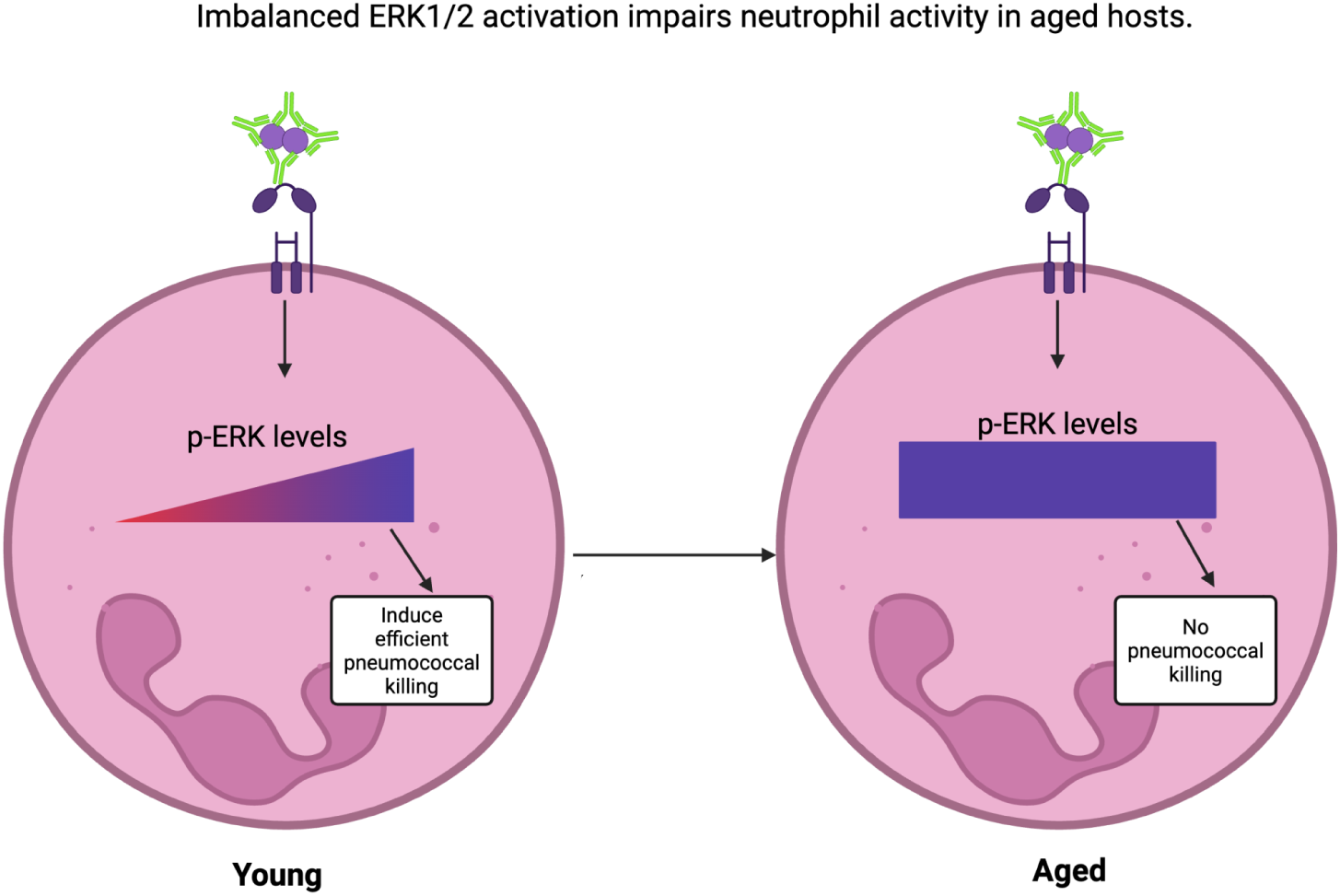

**Graphical Abstract:** Graphic Abstract Created in BioRender. Bou Ghanem, E. (2025) https://BioRender.com/jjoodsu.

## INTRODUCTION

Despite the availability of several licensed vaccines on the market, *Streptococcus pneumoniae* (pneumococcus) infections remain a leading cause of community acquired pneumonia in older adults over the age of 65 (1). Neutrophils, also known as polymorphonuclear leukocytes (PMNs), are cells of the innate immune system that are essential for host protection against infections with *S. pneumoniae*. PMNs are among the first cells recruited to the lungs following pneumococcal infection and previous studies have shown that PMNs are essential for protection not only in naïve hosts, but also in vaccinated hosts (2–5). However, PMN function declines with host aging (3, 6, 7). Prior work from our lab showed that impaired PMN function contributes to the decline in pneumococcal vaccine efficacy seen with aging (3). We observed a decline in PMN function with age in response to *S. pneumoniae* opsonized with sera from hosts immunized with the pneumococcal conjugate vaccine (3). This decline in function persisted when the sera were isolated from young mice indicating that this age-driven defect was intrinsic to PMN function (3). Importantly, adoptive transfer of PMNs from young mice into aged, vaccinated mice prior to pneumococcal infection enhanced vaccine-mediated protection in these mice lowering clinical signs of disease and bacterial burden in the lung (3). Therefore, enhancing PMN function in vaccinated, aged hosts could improve vaccine efficacy, however the signaling pathways controlling PMN function in vaccinated hosts remain incompletely defined.

Activation of complement and Fc receptors on PMNs via complement and antibody mediated opsonophagocytosis is important for host protection during pneumococcal infection (8–11). A major virulence factor contributing to *S. pneumoniae* pathogenesis is the expression of a polysaccharide capsule on the bacterial surface. This capsule helps *S. pneumoniae* evade phagocytosis by immune cells through repulsion as both the capsule and PMNs have an overall net negative charge (12, 13). Deposition of opsonins, in the form of complement and pneumococcal capsular polysaccharide specific antibodies, on the bacterial surface helps promote phagocytosis by neutrophils. Previous studies have shown that certain pneumococcal serotypes can impair complement binding to the capsule surface highlighting the importance of antibody-mediated opsonophagocytosis by PMNs to promote host protection from disease (14, 15). Indeed, all currently licensed vaccines target the pneumococcal polysaccharides and elicit antibodies that enhance bacterial clearance by PMNs (16). In humans, sera opsonic capacity is used as a measure of vaccine immunogenicity (17, 18).

Understanding the pathways regulating PMN function in a vaccinated host is necessary because activation of PMNs by complement receptors and Fc receptors have been shown to trigger distinct signaling pathways, differing PMN antimicrobial functions, and receptor specific changes in gene expression (19, 20). In fact, while complement-mediated bacterial killing by PMNs from aged mice is reduced, these cells are completely unable to kill antibody opsonized bacteria, suggesting larger age-driven decline in PMN responses downstream Fc receptors activation. A better understanding of downstream signaling pathways and how they are altered with host aging is needed to enhance PMN responses in aged, vaccinated hosts.

Mitogen-activated protein kinases (MAPK) are kinases involved in several signaling pathways in response to activation by cell surface receptors. MAPK signaling cascades include extracellular signal regulated kinase (ERK) family, c-Jun N terminal kinase family (JNK), and the p38 kinase family (21, 22). These signaling cascades are activated downstream of many receptor types including growth factor receptors, toll like receptors, GPCRs, and cytokine receptors (21, 22). Additionally, MAPK signaling components are activated downstream of FcγR activation (23–25). Previous work has shown that MAPK signaling regulates PMN responses to infection (26–28). It has also been shown that there is dysregulation in this pathway with host aging (29–31). Prior work from our lab found increased expression of genes associated with MAPK signaling in PMNs isolated from unvaccinated old mice in response to *S. pneumoniae* compared to young controls (30). This increase was associated with increased JNK/AP-1 activation and inhibition of this pathway reversed the age-related defect in killing of complement opsonized *S. pneumoniae* by PMNs (30).

In this study we explored age-driven changes in MAPK signaling in response to complement versus antibody opsonized *S. pneumoniae.* We found age and opsonin dependent changes in MAPK activation in PMNs. We found that aging was associated with heightened levels of basal activation of MAPK pathway components that prevented further activation downstream of Fcγ receptor signaling. Importantly, we identified that balanced activation of ERK1/2 to be crucial for PMN antimicrobial activities against antibody opsonized bacteria following vaccination in both mice and humans.

## MATERIALS AND METHODS

### Mice

Young (2-3 months) and old (20-22 month) C57BL/6 (B6) male mice were purchased from Jackson Laboratories or the National Institute on Aging. All mice were housed in specific pathogen free housing for a minimum of two weeks before use in experiments. All animal studies were performed in accordance with the guidelines in the Guide for the Care and Use of Laboratory Animals and in accordance with the University at Buffalo Institutional Animal Care and Use Committee guidelines.

### Bacteria

*Streptococcus pneumoniae* TIGR4 (serotype 4) was a generous gift from Andrew Camilli (32). Bacteria were grown in Todd Hewitt broth supplemented with 0.5% yeast extract and oxyrase at 37⁰C at 5% CO2 to mid-exponential phase as previously described (33).

### Murine PMN isolation

To isolate PMNs, bone marrow (BM) was flushed from the hind tibias and femurs using RPMI with 10% fetal bovine sera and 2mM EDTA. This cell suspension was strained using 100μm cell strainers. Following lysis of red blood cells, cells were washed and resuspended in PBS. Histopaque 1119 and 1077 were used to separate PMNs via density centrifugation as previously described (34). PMNs were resuspended in Hank’s Balanced Salt Solution (HBSS) 0.1% gelatin with no Ca^2+^ or Mg^2^ at the indicated concentrations prior to use in each assay.

### Generation of immune sera

Young mice were vaccinated with the pneumococcal conjugate vaccine (PCV) Prevnar-13^®^ (Wyeth pharmaceuticals) or mock treated with PBS injected intramuscularly into the caudal thigh muscle. Following vaccination, mice were housed in our facility for four weeks. Mice were then euthanized, and blood was collected by cutting the portal vein. Blood was then centrifuged and sera from each mouse group were pooled and stored at -80⁰C until use. Prior to use in assays, the sera were heat inactivated (HI) at 56⁰C for 40 minutes to inactivate complement proteins in the sera (35). Previously our group found that heat inactivation of sera prevented complement deposition on the pneumococcal surface but did not affect antibody binding to the bacteria (3).

### FcγR Flow Cytometry

PMNs were isolated from the bone marrow of young or old B6 mice. PMNs were infected at a multiplicity of infection (MOI) of 2 with *S. pneumoniae* TIGR4 opsonized with 3% immune sera, or mock infected with 3% sera and buffer alone, for 30 minutes at 37⁰C. Following infection, PMNs were stained with the following antibodies purchased from Invitrogen: anti-Ly6G (1A8), anti-CD16 (002), anti-CD16-2 (012), and anti-CD32b (AT130-2). Cells were then fixed using CytoFix buffer. Fluorescent intensity was measured using BD Celesta and a minimum of 20,000 events were analyzed using FloJo software.

### Phosphorylated ERK1/2 Flow Cytometry

PMNs isolated from the BM of young and old mice were infected with *S. pneumoniae* TIGR4, pre-opsonized with 3% HI immune sera at MOI of 2 for 5 minutes at 37⁰C. Following infection, reactions were fixed using CytoFix buffer and permeabilized with ice-cold 90% methanol. PMNs were then treated with Fc block (anti-mouse clone 2.4G2) and then stained for phosphorylated ERK1/2 (Thr980) (16F8; Cell Signaling), total ERK1/2 (C33E10; Cell Signaling) and Ly6G (1A8; Invitrogen) and secondary antibody anti-Rabbit IgG (12473981; Invitrogen). Percentage of total and phosphorylated ERK1/2 positive cells was determined by flow cytometry using BD Fortessa and a minimum of 20,000 events were analyzed using FloJo software.

### MAPK Phosphorylation

PMNs were isolated from the BM of young and old B6 mice. PMNs were either mock treated or infected with *S. pneumoniae* pre-opsonized with 3% naïve, or HI immune sera at MOI of 2 for 5 minutes at 37⁰C. Following infection PMNs were pelleted and lysed and cell lysates were saved at -80⁰C until use. Cell lysates were analyzed by MAPK phosphorylation array (RayBiotech) per manufacturer’s protocol. Briefly, lysates were incubated on antibody coated membrane specific for 17 different phosphorylated MAPK components. Chemiluminescence of blots was determined using ChemiDoc imaging machine (Bio-Rad) and pixel density was measured using FIJI (ImageJ) software. Positive controls were used to normalize pixel density between blots. Fold changes were calculated from uninfected baselines. Twenty percent differences corresponding to fold changes greater than 1.2 were considered upregulated and below 0.8 were considered down regulated.

### Real Time Quantitative PCR (qPCR)

Bone marrow PMNs isolated from young or old B6 mice were treated with specific ERK1/2 inhibitor PD98059 (50μM) or vehicle control for 5 minutes at room temperature immediately before infection. PMNs were then infected with *S. pneumoniae* TIGR4 pre-opsonized with 3% HI immune sera at MOI 2 at 37⁰C for 5 minutes. Following infection, the cell pellet was collected via centrifugation and RNA was extracted and isolated by Qiagen RNeasy Mini kit per manufacturer protocol. 500ng of purified RNA was converted to cDNA using SuperScript VILO^TM^ cDNA synthesis kit (Life Technologies, USA) according to the manufacturer’s protocol. cDNA was stored at -20⁰C until use. qPCR was performed using an Applied Biosystems® ViiA-7 real time PCR system (Life Technologies, USA). GAPDH gene expression was assessed as a control. TaqMan probes from Life Technologies used include: 1. Mm00457274_g1 (DUSP1 gene) and 2. Mm99999916_g1 (GAPDH gene). Data were analyzed by the comparative threshold cycle (CT) method, normalizing the CT values for target gene expression to those for GAPDH. Relative quantity (RQ) values were calculated by the formula RQ =2^-ΔΔCT^. To obtain the relative expression values, RQ values in each experiment were normalized to the young, uninfected, vehicle control condition.

### Opsonophagocytic killing assay (OPH)

Bone marrow PMNs from young or old B6 mice were treated with specific ERK1/2 inhibitor PD98059 (CellSignalling) (50μM) or vehicle control for 5 minutes at room temperature immediately before infection. PMNs were then infected with *S. pneumoniae* TIGR4 pre-opsonized with 3% naïve or HI immune sera at MOI 0.01 at 37⁰C for 40 minutes. Reactions were then stopped and placed on ice for 2 minutes. Following this incubation reactions were plated on blood agar to enumerate bacterial CFU. Pneumococcal killing percentage was determined with respect to no PMN control wells treated with the same conditions.

### Annexin PI Staining

PMNs isolated from the BM of young mice were pretreated with either vehicle control or specific ERK1/2 inhibitor PD98059 (50μM) for 30 minutes at 37⁰C. Following treatment, PMNs were mock infected with sera alone or infected at MOI 2 with *S. pneumoniae* pre-opsonized with 3% naïve sera for 40 minutes. Following infection, reactions were stained for viability using Annexin-PI staining and flow cytometry was used to determine the percentage of live, apoptotic, and necrotic cells. BD Fortessa was used and a minimum of 20,000 events were analyzed using FloJo software.

### Human participants

All work with human donors was approved by the University at Buffalo Human investigation Review Board (IRB). Young adult healthy human donors (21-40 years old, male, and female) with no prior history of pneumococcal immunization were recruited and individuals that were taking medications, were pregnant or had chronic or acute infections were excluded from the study. All enrolled donors signed approved informed consent forms. Donors were administered the pneumococcal conjugate vaccine Prevnar-20^®^ via intramuscular injection as per manufacturer’s instructions. Blood was drawn prior to and at week four following vaccination. Blood draws occurred in the morning between 9 and 10 AM.

### PMN and sera isolation from human donors

Whole blood samples were collected using acid citrate/dextrose as an anticoagulant. The samples were centrifuged, and cell free sera collected. PMNs were then isolated via negative selection using the EasySep^TM^ Direct neutrophil isolation kit (StemCell) per manufacturer protocol. PMNs were resuspended finally in Hank’s Balanced Salt Solution (HBSS) 0.1% gelatin with no Ca^2+^ or Mg^2+^ prior to use in opsonophagocytic killing assay.

### Statistics

Statistics were analyzed using Prism 9 (GraphPad). All data are shown as mean +/- the standard deviation. Normality of data was tested via Shapiro-Wilk test prior to statistical analysis. Significant difference was determined by One Sample t and Wilcoxon test, or unpaired student’s t test as indicated. *p <0.05* was considered significantly different and is indicated in the figures with *.

## RESULTS

### Differential activation of PMN MAPK pathway components by complement opsonized versus antibody opsonized *S. pneumoniae*

As the age-related decline in PMN antimicrobial activity was significantly worsened when PMNs were infected with antibody opsonized versus complement opsonized bacteria (3), we analyzed how signaling downstream of both complement and Fcγ receptor activation changes in PMNs. Using a MAPK pathway phosphorylation array (RayBiotech), we first analyzed how phosphorylation of different MAPK pathway components change in PMNs isolated from young mice at baseline and following infection with *S. pneumoniae* opsonized with sera from age-matched naïve mice or heat inactivated (HI) sera from mice immunized with the pneumococcal conjugate vaccine (PCV) to stimulate complement-mediated and antibody-mediated PMN responses respectively (**Figure 1A**). Fold change from uninfected basal levels were calculated for naïve and HI immune conditions for all 17 components tested (**Figure 1B**). Using a 20% change (1.2-fold change) as the cutoff, we found that most changes that occurred were common between the infection conditions, while a few were specific to each opsonin. Several proteins showed increased phosphorylation above baseline in both conditions including P70S6K, RSK2, MKK6, MSK2, ERK1/2, and mTOR, while RSK1 phosphorylation was the only one that displayed a decrease in both infection conditions (**Figure 1B and C**). ERK1/2 and mTOR phosphorylation were highly upregulated upon infection with ERK1/2 phosphorylation increasing 16-fold and mTOR phosphorylation increasing between 3 and 4-fold (**Figure 1B**). The three proteins that were only phosphorylated above baseline when PMNs were infected with pneumococcus opsonized with complement were MEK, CREB, and GSK3a (**Figure 1B and C**). In contrast, GSK3b and p38 increased in phosphorylation only upon infection with pneumococcus opsonized with antibodies (**Figure 1B and C**). These data suggest that complement-mediated, and antibody-mediated activation of PMNs following infection activate several MAPK pathway components differently, supporting previous studies (36).

**Figure 1:**
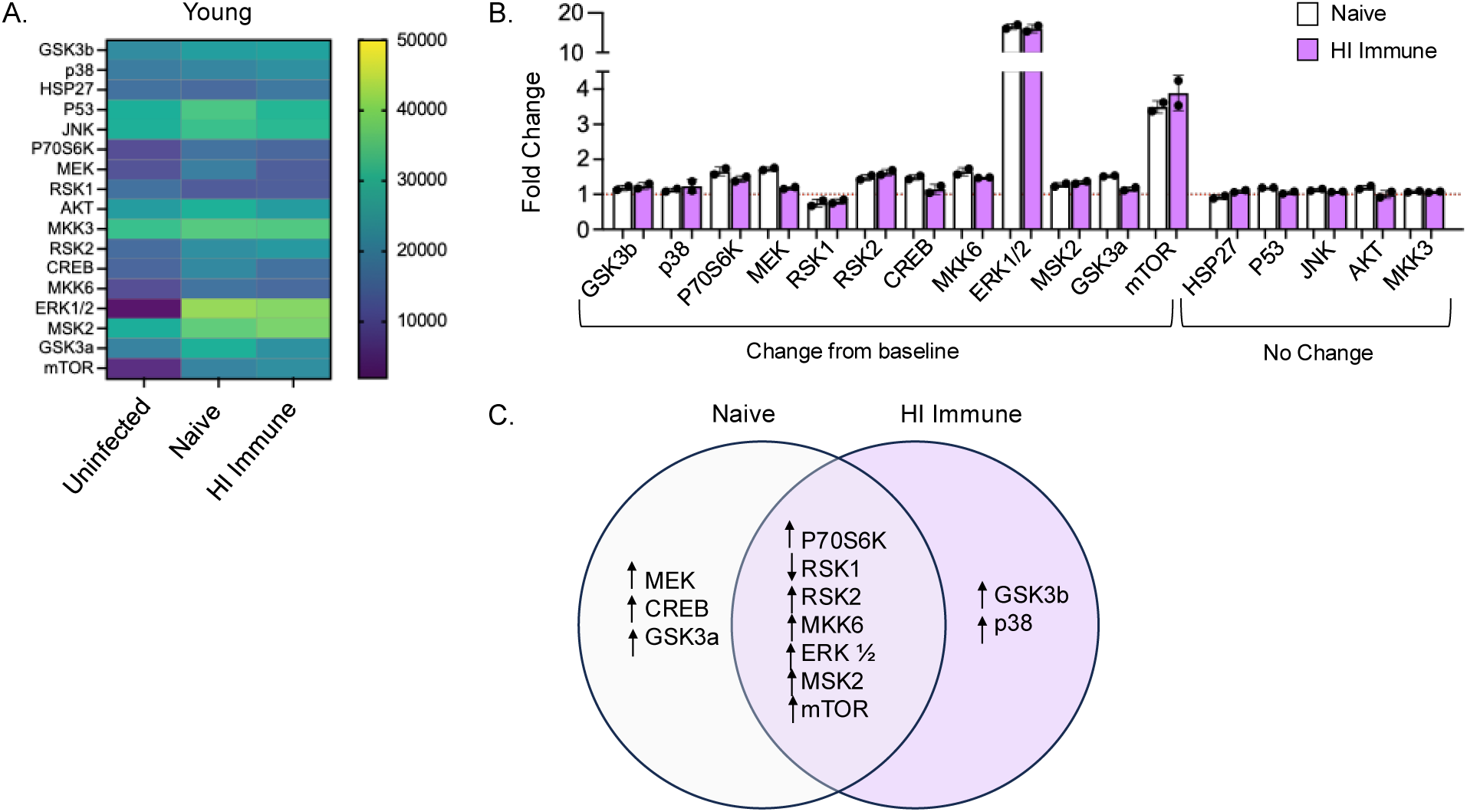
Changes in PMN MAPK pathway activation in young mice. PMNs were isolated from the BM of young C57BL/6 mice and infected with *S. pneumoniae* opsonized with naïve or HI immune sera. MAPK phosphorylation was determined by MAPK phosphorylation array. (A) Heat map showing phosphorylation levels in uninfected and infected PMNs. (B) Fold changes upon infection for naïve (white) and HI immune (purple) compared to uninfected baseline. Twenty percent difference was set as a cut-off where fold changes greater than 1.2 were considered increased above baseline and fold changes less than 0.8 were considered decreased from baseline. Fold changes between 0.8 and 1.2 were considered to be no change from baseline expression. (C) Venn diagram indicating shared and separate MAPK component changes in both naïve and HI immune infection conditions. Up arrows indicate fold change greater than 1.2 and down arrows indicate a fold change less than 0.8. Lysates were pooled from n=3 biological replicates and samples were run in duplicates.

### PMNs from aged hosts display less activation of MAPK pathway components in response to antibody opsonized compared to complement opsonized *S. pneumoniae*

To better understand the age-related changes in PMN signaling following FcγR activation, we next asked does MAPK component phosphorylation change following infection in aged hosts. PMNs were isolated from aged mice and protein phosphorylation was measured at uninfected baseline and upon infection with *S. pneumoniae* opsonized with sera from naïve mice or heat inactivated (HI) sera from immunized mice (**Figure 2A**). To probe the intrinsic changes in PMN activity, independent of the confounding effects of the age-driven decline in antibody responses to immunization (6, 17), sera used were from young mice. Fold change from uninfected basal phosphorylation levels were calculated to determine activation with 1.2-fold change used as a cutoff (**Figure 2B**). Unlike what we found in PMNs from young mice, all increases in MAPK component phosphorylation in PMNs from aged mice occurred upon infection with complement but not antibody opsonized pneumococci. An increase in phosphorylation above baseline was observed in JNK, P70S6K, AKT, MKK3, RSK2, CREB, MKK6, GSK3a, and mTOR (**Figure 2B and C**). In contrast, challenge with antibody opsonized bacteria did not result in any increase in MAPK component phosphorylation in PMNs from aged mice (**Figure 2B and C**). In fact, decreased phosphorylation of GSK3b and p38 were the only changes in phosphorylation of the pathway components tested in response to infection with antibody opsonized *S. pneumoniae* by PMNs from aged mice (**Figure 2B and C**). This is opposite of what was found in young mice, where GSK3b and p38 phosphorylation increased following PMN infection with antibody opsonized bacteria (**Figure 1C**). These data suggest that activating complement receptors on PMNs from aged hosts results in changes in the MAPK pathway similar to young hosts. In contrast, activation of the MAPK pathway downstream of Fcγ receptor signaling is impaired with age.

**Figure 2:**
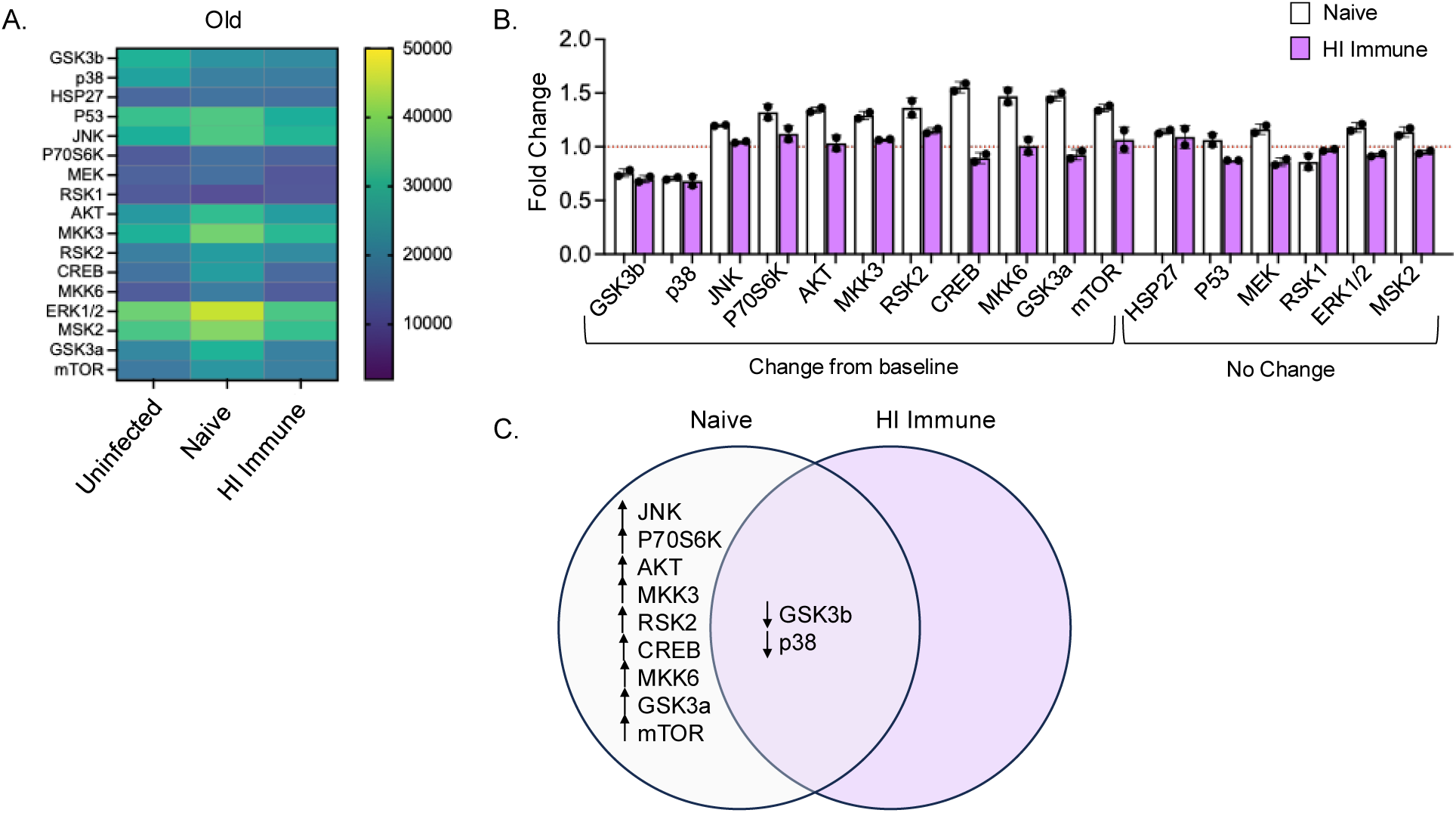
Changes in PMN MAPK pathway activation in old mice. PMNs were isolated from the BM of old C57BL/6 mice and infected with *S. pneumoniae* opsonized with naïve or HI immune sera. MAPK phosphorylation was determined by MAPK phosphorylation array. (A) Heat map showing phosphorylation levels in uninfected and infected PMNs. (B) Fold changes upon infection for naïve (white) and HI immune (purple) compared to uninfected baseline. Fold changes greater than 1.2 were considered increased above baseline and fold changes less than 0.8 were considered decreased from baseline. Fold changes between 0.8 and 1.2 were considered to be no change from baseline expression. (C) Venn diagram indicating shared and separate MAPK component changes in both naïve and HI immune infection conditions. Up arrows indicate fold change greater than 1.2 and down arrows indicate a fold change less than 0.8. Lysates were pooled from n=3 biological replicates and samples were run in duplicates.

When we overlaid the data on the MAPK signaling cascade downstream of several stimuli, we found that age resulted in lack of increased phosphorylation of MAPKKs, MAPKs and effector molecules (**Figure 3A**) in response to acute stimulation with antibody opsonized bacteria. We found that the MAPK cascades activated in PMNs from young mice are also found downstream of GPCR, and inflammatory cytokine receptors while none of these pathways were activated with age (**Figure 3A**). In contrast, in response to acute stimulation with naïve sera opsonized bacteria, pathways activated in young hosts are also activated in old hosts downstream of several receptor types (**Figure 3B**). These data suggest there is an age-related defect in MAPK signaling downstream of several different receptor types when PMNs encounter *S. pneumoniae* in vaccinated hosts.

**Figure 3:**
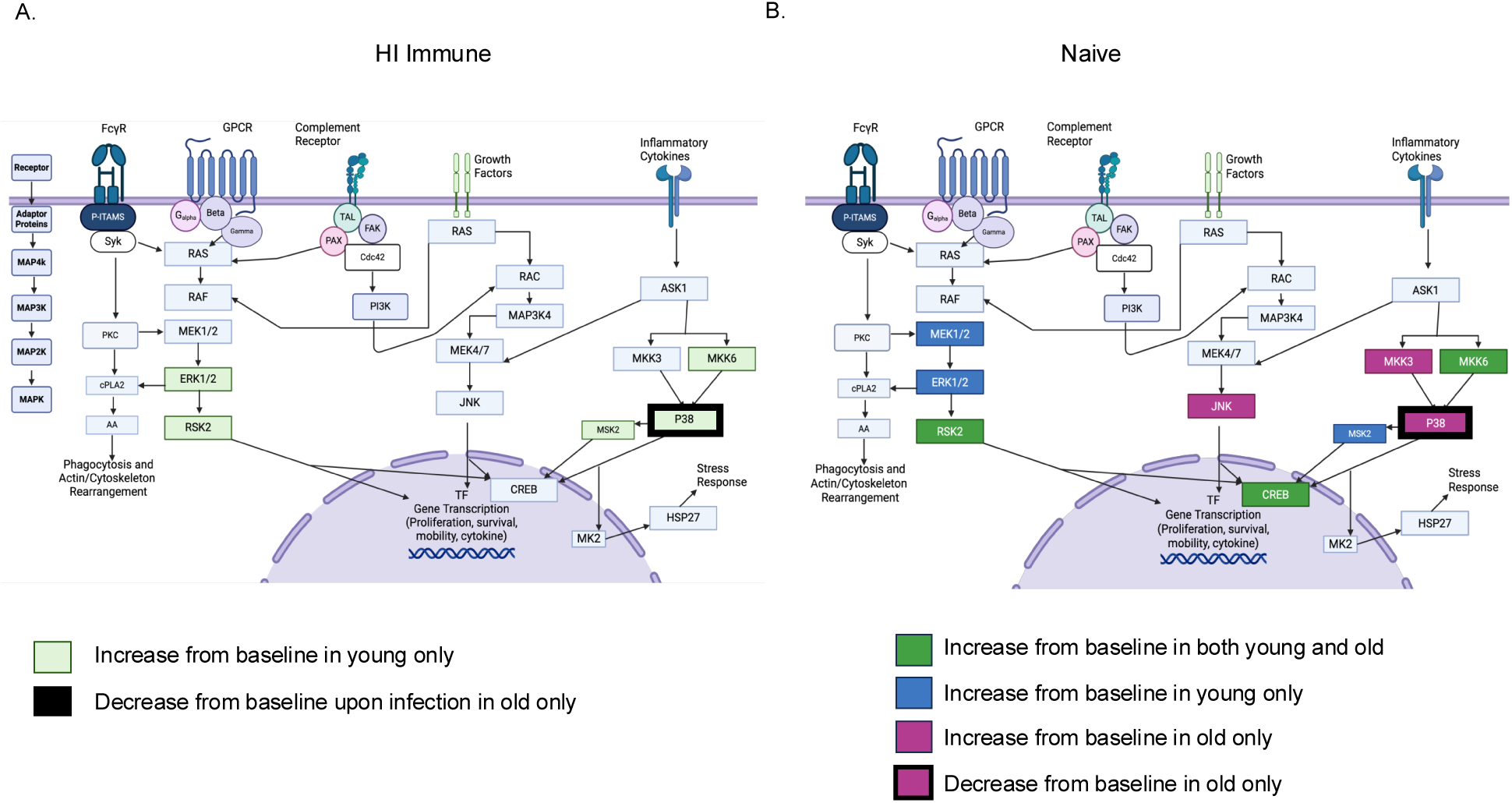
MAPK signaling pathways following infection. General MAPK signaling pathways following integrin receptor, Fc receptor, G-protein coupled receptor, growth factor receptor, and cytokine receptor activation. Changes in MAPK pathway component phosphorylation from baseline upon infection with *S. pneumoniae* opsonized with (A) HI immune sera and (B) naïve sera in PMNs from young and old C57BL/6 mice are overlayed on the pathway (23, 24, 38–41, 63). Created in BioRender. (A) Bou Ghanem, E. (2025) https://BioRender.com/wn94z0h and (B) Bou Ghanem, E. (2025) https://BioRender.com/wn94z0h.

### Age-driven changes in FcγR expression on PMNs do not account for reduced activation of MAPK pathway components in response to antibody opsonized bacteria

As PMNs were refractory to activation of Fcγ receptors, we asked if this is caused by changes in FcγR expression on the surface of PMNs such as decline in activating receptors and increase in inhibitory ones. We asked if FcγR surface expression changes with host aging by measuring receptor expression on the surface of PMNs isolated from young and aged mice by flow cytometry (gating strategy, **Supplemental Figure 1A**). Compared to young controls, we found that there was a significant increase in the percentage of PMNs expressing FcγRIII (CD16-activating) both at baseline and upon infection with *S. pneumoniae* when isolated from aged mice (**Figure 4A**). In contrast, we found that with age there was no significant difference in the percent of cells expressing of FcγRIIb (CD32b-inhibitory) or FcγRIV (CD16-2-activating) (**Supplemental Figure 1B** and **Figure 4A**). When examining the level of receptor expression (gMFI) we found no difference in any of the Fcγ receptors across host age (**Supplemental Figure 1C** and **Figure 4B**. These data show that aging is not associated with reduced Fcγ receptor expression on PMNs.

**Figure 4:**
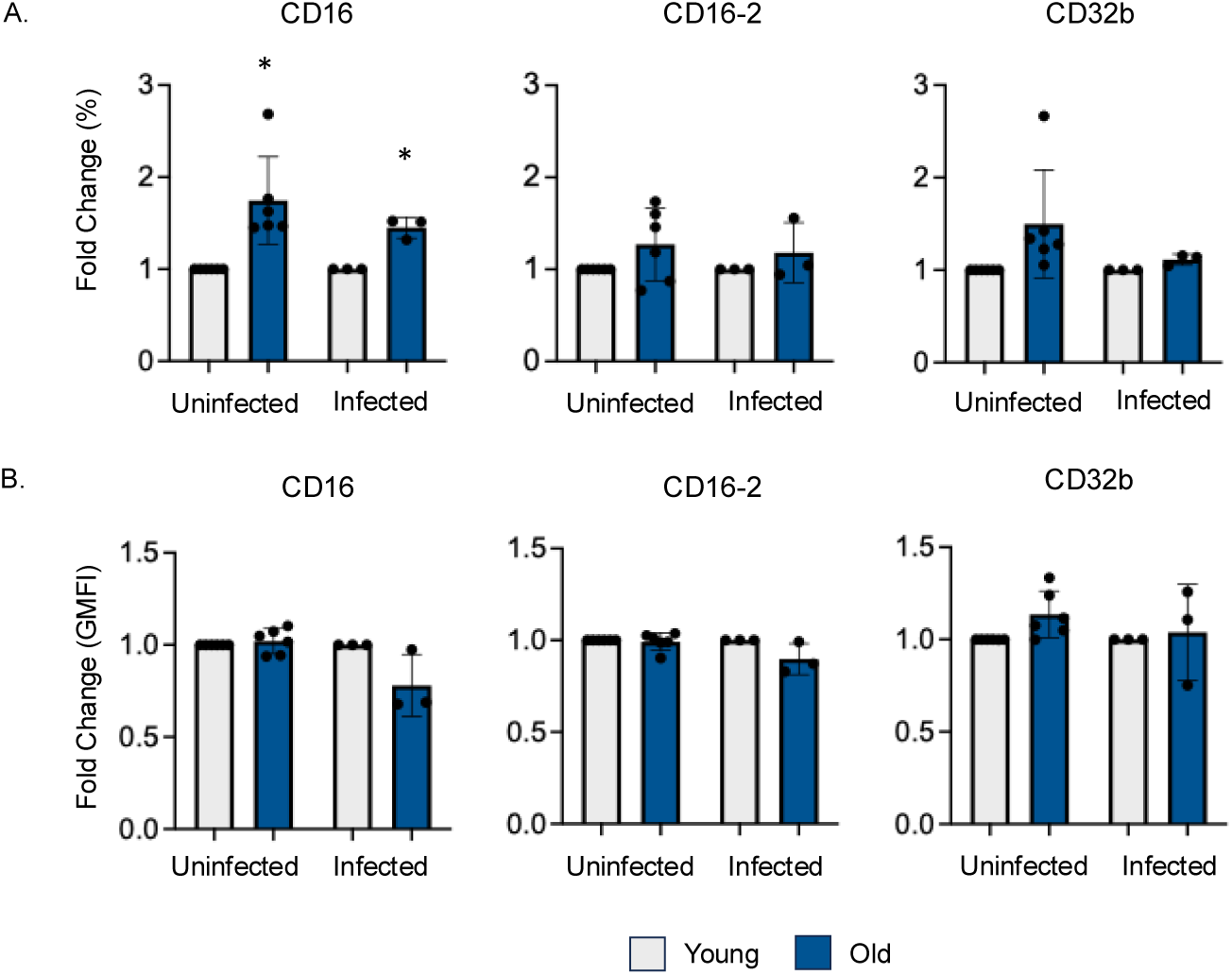
Change in FcγR expression with age. Surface expression of FcγRs CD16, CD16-2 and CD32b on BM PMNs from young and old C57BL/6 mice at uninfected baseline and following infection with *S. pneumoniae* opsonized with immune sera. The fold changes for (A) % expression and (G) geometric mean florescent intensity (GMFI) were calculated from young controls and pooled from separate mice, n=6 for uninfected and n=3 infected. * indicates *p<0.05* as determined one sample t and Wilcoxon test.

### Basal activation of MAPK components renders PMNs from aged hosts hyporesponsive to acute stimulation with antibody opsonized pneumococci

To better understand the age-related changes in MAPK pathway activation, we compared phosphorylation levels of each of the MAPK pathway components in PMNs from young and aged mice at uninfected baseline (**Figure 5A**), and upon infection with complement (**Figure 5B**) or antibody opsonized (**Figure 5C**) *S. pneumoniae*. At baseline there was a significant increase in basal activation of several components in PMNs from aged hosts compared to young controls (**Figure 5A**). This included increased basal phosphorylation of ERK1/2, mTOR, p38, GSK3b, P70S6K, MEK, RSK2, MKK6, and MSK2. In contrast, upon infection many proteins had similar phosphorylation levels in PMNs from young and aged mice in both the naïve and immune infection conditions.

**Figure 5:**
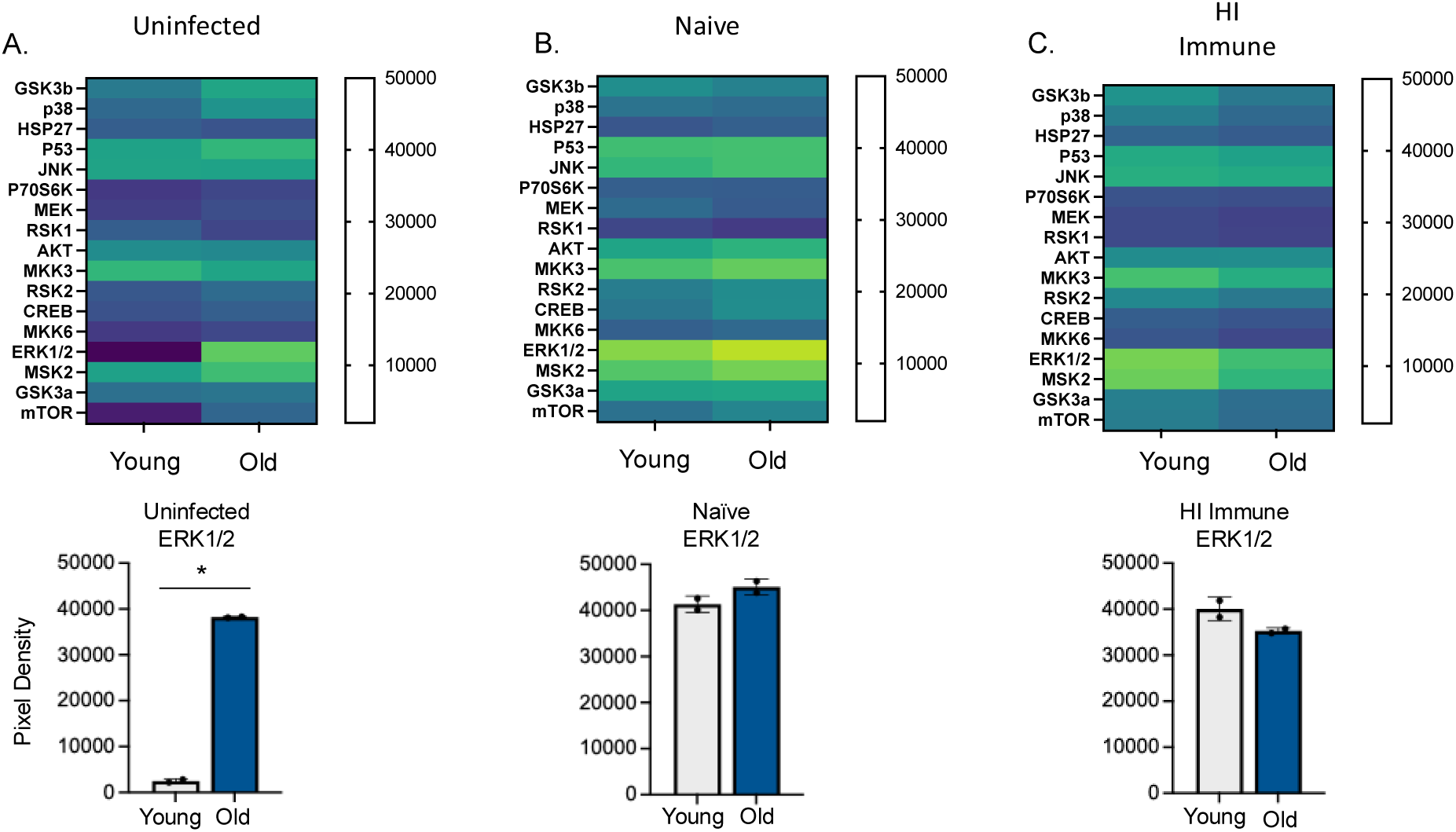
Basal MAPK phosphorylation changes with host aging. PMNs isolated from the BM of C57BL/6 young or old were infected with *S. pneumoniae* opsonized with naïve or HI immune sera. Phosphorylation levels of MAPK components are compared between PMNs from young and old mice at uninfected baseline (A), and upon infection with pneumococcus opsonized with naïve (B), or HI immune (C) sera. Phosphorylation levels of ERK1/2 are quantified below heat maps. Lysates were pooled from n=3 biological replicates and samples were run in duplicates.

Of particular interest was ERK1/2 where we observed 15-fold higher basal levels of ERK1/2 phosphorylation in PMNs from aged hosts compared to young controls. In contrast, while ERK1/2 phosphorylation was increased about 16-fold from baseline in response to all infection conditions in young (**Figure 1B**) mice, no increase was observed in aged mice (**Figure 2B**). These findings suggest that ERK1/2 is already activated in aged mice prior to infection (**Figure 5A**) and that this pathway is hyporesponsive to further stimulation. It is possible that the already elevated levels of ERK1/2 at baseline impair the ability of neutrophils to respond to pneumococcal infection if the pathway components cannot be stimulated further.

### Inhibition of ERK1/2 reverses the age-driven defect in killing of antibody opsonized *S. pneumoniae* by neutrophils

We next asked if basal activation of ERK1/2 with age impairs PMN antibacterial activity. To test that, we treated PMNs with the specific ERK1/2 inhibitor PD98059. We confirmed that ERK1/2 inhibition had no effect on the viability of PMNs using Annexin-PI staining (**Supplemental Figure 2A**). The ability of PD98059 to inhibit ERK1/2 was confirmed by measuring phosphorylation by flow cytometry (**Supplemental Figure 2B and C**). We then performed an opsonophagocytic killing assay to measure bacterial killing by PMNs. We found that inhibition of ERK1/2 signaling had no effect on the percentage of *S. pneumoniae* killed by PMNs from young or aged hosts when the bacteria were opsonized with sera from naïve hosts (**Figure 6A**). These finding suggest that ERK1/2 does not play a role in complement-mediated uptake and killing of pneumococci by PMNs.

**Figure 6:**
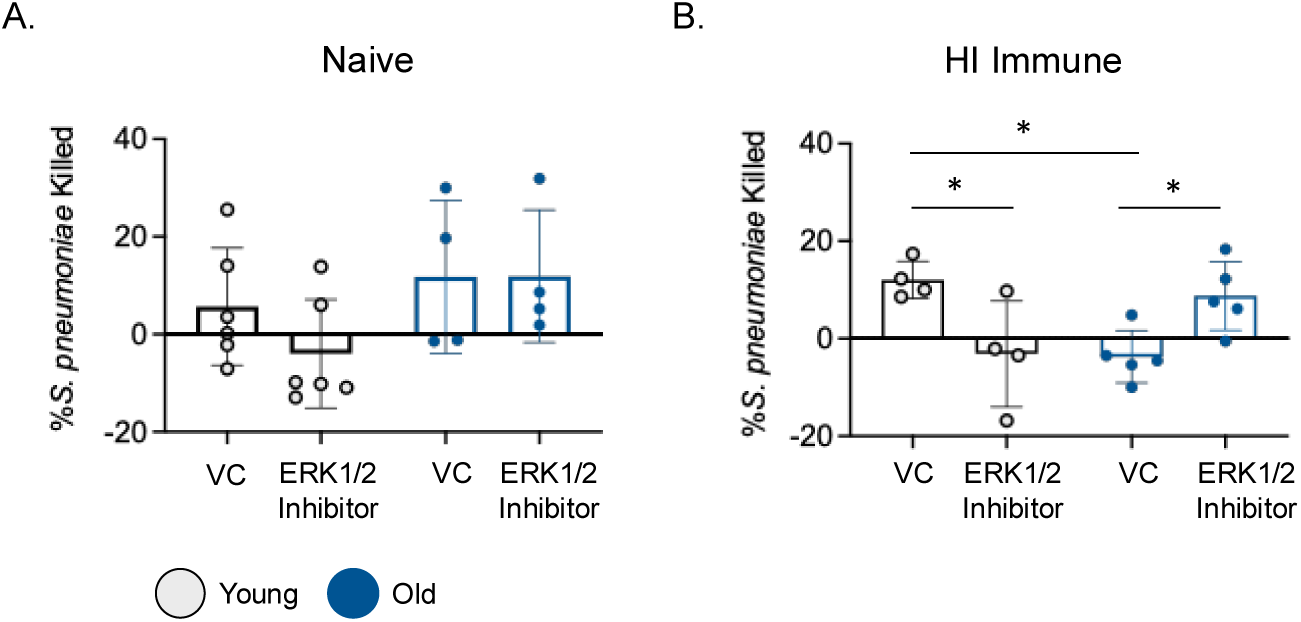
ERK1/2 inhibition affects PMN antimicrobial function. PMNs were isolated from the BM of young or old C57BL/6 mice. PMNs were then infected with *S. pneumoniae* pre-opsonized with naïve (A) or HI Immune (B) sera. The percentage of bacteria killed was then calculated in comparison to a no PMN control under the same treatment conditions. Data are pooled from separate experiments and each dot represents a separate mouse (biological replicate). * indicates *p<0.05* as determined by student’s unpaired t test.

When we tested the role of ERK1/2 on antibody-mediated responses, we found a significant decrease in the ability of PMNs isolated from young mice to kill *S. pneumoniae* opsonized with HI sera from immune hosts following ERK1/2 inhibition (**Figure 6B**). These data suggest that in young mice, ERK1/2 activation upon infection is required for efficient bacterial killing following FcγR activation of PMNs. In contrast, ERK1/2 inhibition of PMNs from aged mice resulted in a significant, 10-fold increase in the ability of PMNs to kill antibody opsonized *S. pneumoniae* (**Figure 6B**). These data suggest that inhibition of basal ERK1/2 activation of PMNs from aged mice rescues their antimicrobial activity. Taken together, these data suggest that balanced activation of the ERK1/2 pathway is required for killing of antibody opsonized bacteria by PMNs and that any aberrant changes in this pathway (over or under activation) impairs PMN antimicrobial function.

### Age-driven changes in dual specificity phosphatases 1 (DUSP-1) expression in PMNs

Phosphatases are important regulators of MAPK signaling. Dual specificity phosphatases (DUSPs) are a family of phosphatases that dephosphorylate threonine and tyrosine residues on MAPKs to inactivate and regulate the duration of signaling (37). To determine if changes in DUSP1 levels were contributing to the age-related increase in basal ERK1/2 phosphorylation, we used qPCR to measure expression of DUSP1 in PMNs treated with vehicle control or specific ERK1/2 inhibitor PD98059 at baseline and upon infection (**Figure 7**). PMNs isolated from young mice had a significant decrease in DUSP1 expression upon pneumococcal infection when compared to the uninfected baseline (**Figure 7**). These data suggest the in PMNs from young mice, infection decreases DUSP1 expression which may contribute to the increase in ERK1/2 phosphorylation upon infection that was observed (**Figure 1B**). In contrast, pneumococcal infection did not reduce DUSP1 expression compared to baseline in PMNs from aged mice and there was a trend towards increased DUSP1 expression in response to infection (**Figure 7**). Therefore, with age, lack of DUSP1 inhibition in response to infection may contribute to lack of further increase in ERK1/2 activation upon acute stimulation (**Figure 7**). In fact, treatment with ERK1/2 inhibitor, which rescued the antimicrobial activity of PMNs from aged hosts against antibody opsonized bacteria (**Figure 6B**), prevented the increase in DUSP1 expression in response to infection. Overall, these data suggest that DUSP1 levels are altered with host age.

**Figure 7:**
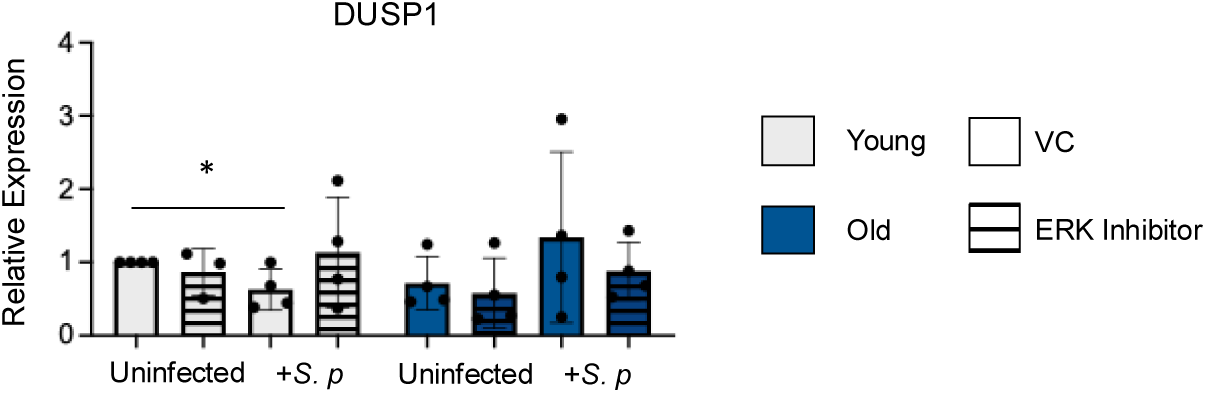
DUSP1 levels in PMNs from young and old mice. BM PMNs from young and old C57BL/6 mice were infected with *S. pneumoniae* opsonized with HI immune sera. Prior to infection PMNs were treated with VC or ERK1/2 inhibitor PD98059. DUSP1 levels were quantified by qPCR at baseline and upon infection. Data are pooled from n=4 biological replicates. * indicates significance (*p<0.05*) as determined by student’s unpaired t test.

### ERK1/2 activation is required for the ability of PMNs from humans immunized with the pneumococcal conjugate vaccine to kill *S. pneumoniae*

Having found that balanced ERK1/2 activation is crucial for PMN activity against antibody opsonized *S. pneumoniae* in mice, we wanted to determine the clinical relevance of these findings in humans. We tested the effect of ERK1/2 inhibition on PMNs isolated from the blood of young human participants before and after administration of the PCV vaccine. PMNs were isolated from the blood of donors at week 0 (prior to vaccination) and week 4 following vaccination. An opsonophagocytic killing assay was used to determine the percentage of *S. pneumoniae* killed by vehicle control and ERK1/2 inhibitor treated PMNs. PMNs were infected with pneumococci opsonized with the donor’s own sera that were heat inactivated to isolate the role of FcγR activation. We found that at week 0, prior to vaccination there was no significant difference in the amount of *S. pneumoniae* killed by PMNs treated with ERK inhibitor when compared to VC (**Figure 8**). This is similar to what was seen previously in murine experiments with no difference in killing in the absence of anti-pneumococcal antibodies. At week 4 following vaccination, there was a significant decrease in the ability of PMNs to kill *S. pneumoniae* when treated with ERK inhibitor compared to VC (**Figure 8**). These data indicate that in young hosts, inhibition of ERK1/2 signaling impairs the ability of human PMNs to kill *S. pneumoniae* in an FcγR-mediated manner.

**Figure 8:**
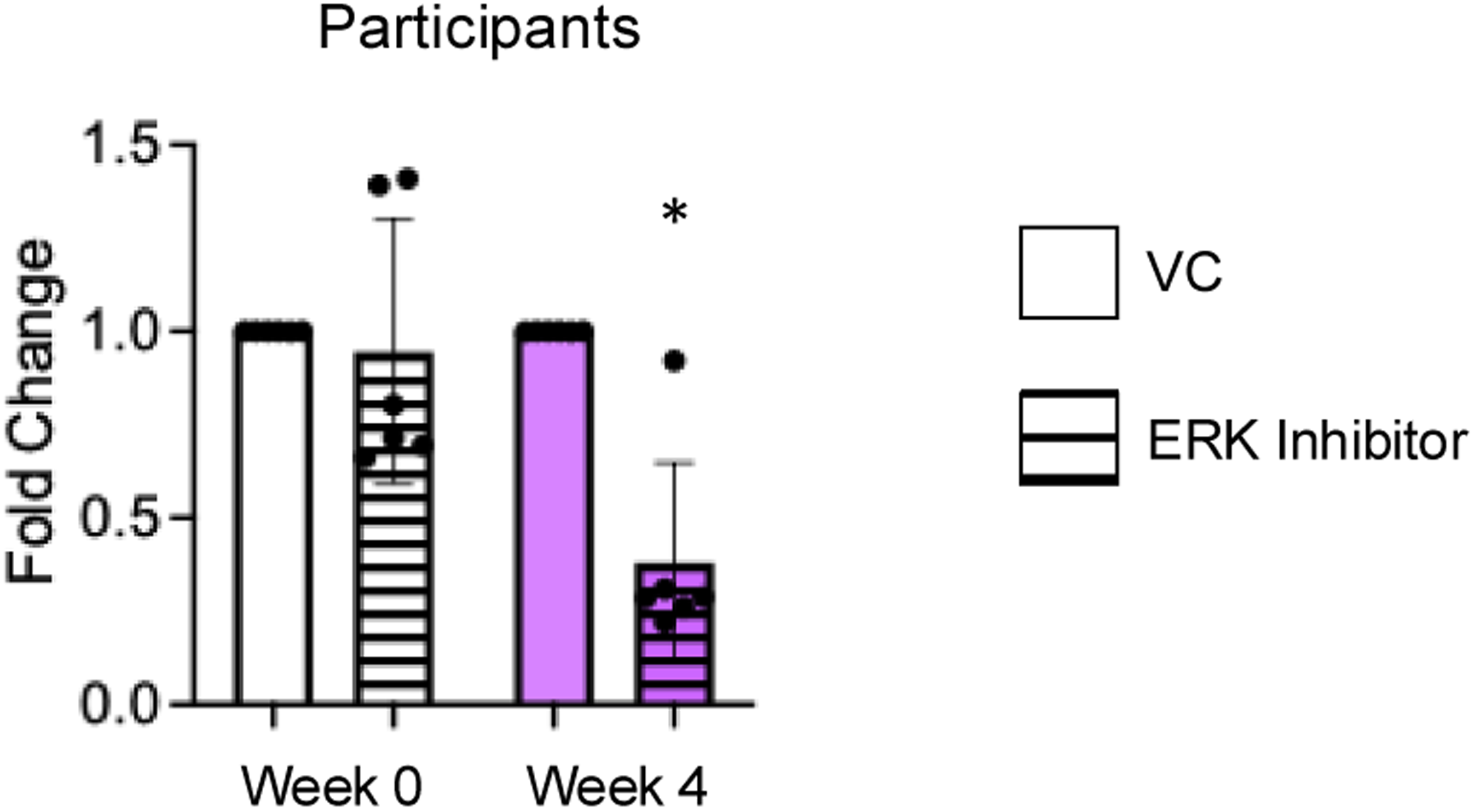
ERK1/2 inhibition decreases the ability of PMNs from young human donors to kill *S. pneumoniae* opsonized with antibodies. PMNs were isolated from the blood of 6 separate young, healthy, volunteers at week 0 and week 4 with respect to vaccination with PCV. Following isolation, PMNs were treated with ERK1/2 inhibitor PD98059 or VC and infected with *S. pneumoniae* opsonized with the donor’s own heat inactivated sera from the same timepoint. Percentage of pneumococcal killing was calculated and killing is shown as fold change from VC. * indicates significantly different from one (*p<0.05)* as determined by One Sample t and Wilcoxon test.

## DISCUSSION

MAPK signaling cascades are important mediators of “outside-in” cell signaling following membrane receptor activation (21, 22). MAPKs respond to mitogens and stress signals to regulate cellular proliferation, and these pathways have also been shown to be important during PMN responses to infection (21, 22, 26, 28). MAPK pathway signaling exhibits cross talk and interaction with several signaling pathways in the cell and for this reason MAPKs are found downstream of several receptor types including Fc receptors and complement receptors (21, 23, 38–41). Previously, we found that the signaling pathways regulating PMN responses to pneumococcal infection differ based on the opsonin activating the PMNs, i.e. complement mediated vs antibody mediated (42). Here, we found that *S. pneumoniae* infection stimulated MAPK pathway activation in PMNs. However, activation of these pathways differed following complement or antibody mediated PMN activation. Additionally, these pathways were altered with host aging. While there were some changes in MAPK component phosphorylation following PMN infection with complement opsonized bacteria with age, infection with antibody opsonized *S. pneumoniae* failed to induce MAPK activation above baseline. This lack of responsiveness was associated with already elevated basal levels in phosphorylation of several components in aged mice including ERK1/2 that resulted in an inability to further respond upon acute stimulation. Importantly, balanced activation of the ERK1/2 pathway was required for killing of antibody opsonized bacteria by PMNs and aberrant changes in this pathway including over or under activation impaired PMN antimicrobial function in murine models and human participants. This study identifies age-driven defects in Fcγ receptor signaling and expands our understanding of the differential role of MAPK ERK1/2 pathway in PMN responses.

One key finding of this study was the elevated basal levels of phosphorylation of several MAPK component in PMNs from aged hosts that rendered the host unable to further respond to acute stimuli. Similar results have been found in prior studies where elevated levels of JNK, p38 or ERK1/2 with age impair PMN function (29–31). Elevated p38 levels in aged hosts resulted in reduced efferocytosis of apoptotic PMNs by macrophages (29). Increased ERK1/2 activation in PMNs from aged hosts impaired the ability of inflammatory signal GM-CSF to delay PMN apoptosis (31). While increased expression of JNK/AP-1 was found to inhibit the ability of PMNs from aged mice to kill complement opsonized pneumococci (30). We found here that basal ERK1/2 phosphorylation was 15-fold higher in PMNs from aged mice when compared to young. However, upon infection in both the naïve and immune conditions, PMNs from young and aged mice had similar ERK1/2 phosphorylation levels suggesting that ERK1/2 cannot be activated further upon infection with age. These data suggest that increased basal MAPK phosphorylation in PMNs from aged mice prevents activation of antimicrobial responses upon acute infection. The reason behind this elevated basal activation is unclear. Aging results in chronic, widespread, low-grade inflammation, known as inflammaging (43–45) driven by increased inflammatory cytokines and proinflammatory mediators. Inflammaging is proposed to contribute to the reduced PMN function with age (45, 46). Previous studies have shown that chronic exposure to inflammation can reduce PMN antimicrobial responses when restimulated with TNF, demonstrating an “exhausted” PMN phenotype (47). The increased basal activation of ERK1/2 with no increase in phosphorylation upon pneumococcal infection may be due to immune tolerance where following chronic repeated exposure to stimuli, immune cells are no longer able to activate gene transcription or perform normal functions in response to acute stimulation (48). Therefore, inflammaging may be activating MAPK pathways in PMNs rendering them unable to respond effectively to infection.

Phosphatases are important regulators of MAPK signaling, DUSP1 dephosphorylates both ERK1/2 and p38 to regulate activation (37, 49). Altered DUSP1 phosphatase expression with host aging has been shown in alveolar macrophages in response to chronic TNF exposure where DUSP1 was elevated which induced a suppression of MAPK signaling and was associated with higher susceptibility to *Streptococcus pneumoniae* infection (38). Here, we found that upon infection, DUSP1 levels significantly decreased in PMNs from young but not aged mice. DUSP1 instead trended toward increased expression in response to infection in aged hosts. It is possible that elevated DUSP1 levels in PMNs from aged mice during pneumococcal infection may contribute to lack of further ERK1/2 phosphorylation and diminished pneumococcal killing. In fact, ERK1/2 inhibition of PMNs from aged mice trended toward lower DUSP1 levels which may explain why ERK inhibition increased pneumococcal killing only in aged mice. These data are supported by the known immunosuppressive role of DUSP1, where DUSP1 knock out macrophages had increased pro-inflammatory cytokine production and glucocorticosteroids used in chronic inflammatory diseases produce immune suppression partly due to activation of DUSP1 (37, 50–52).

ERK1/2 activation regulates PMN antimicrobial responses in several models of infection and in response to stimulation by bacterial products. These PMN responses include PMN adhesion, NETosis formation, granule mobilization to engulfed pathogens, phagocytosis, delayed apoptosis, chemotaxis, and respiratory burst (26–28, 36, 53–55). Here we found that balanced activation of the ERK1/2 pathway was required for killing of antibody opsonized bacteria by PMNs and over or under activation impaired PMN antimicrobial function. While inhibition of ERK1/2 significantly reduced pneumococcal killing in young hosts, it rescued the age-related defect in pneumococcal killing. We hypothesize that the opposing effects of ERK1/2 inhibition on PMN response to infection by PMNs isolated from young and aged mice is due to the increased basal activation (described above). PMNs from aged mice have higher basal activation that is refractory to further stimulation from infection, resulting in diminished antimicrobial response. It is possible that ERK1/2 inhibition in PMNs from aged mice prior to infection enhances ERK1/2 activation in response to acute stimulation, when the PMNs encounter the pneumococcus, enabling a more robust antimicrobial response. Importantly, we verified this in human participants and showed that activation of ERK1/2 was required for the ability of PMNs to kill antibody opsonized bacteria in donors vaccinated with PCV. These data suggest that ERK1/2 is a clinically relevant pathway regulating PMN responses in vaccinated hosts.

The effect of ERK1/2 in PMNs in both mice and humans was only found in the presence of antibodies in the sera against *S. pneumoniae* but not in response to complement opsonized bacteria. In prior work we found that other MAPK components, in particular JNK/AP-1 was required for killing of complement opsonized bacteria and that elevated basal levels of this pathway impaired the ability of PMNs from aged mice to kill bacteria (30). These findings demonstrate that MAPK components have differential roles downstream Fcγ versus complement receptor signaling. Further studies are needed to identify how ERK1/2 signaling is mediating killing of *S. pneumoniae* and to identify why this pathway is specific to antibody mediated PMN activation.

In comparing Fcγ versus complement receptor activation in young hosts, there were many commonalities in MAPK signaling activation, however we found that GSK3b and p38 were only phosphorylated in response to antibody opsonized *S. pneumoniae.* Interestingly, this was the only pathway that changed in aged hosts in response to antibody opsonized *S. pneumoniae,* where decreased phosphorylation of GSK3b and p38 was observed. GSK3b (glycogen synthase kinase 3b) is a known regulator of inflammation following bacterial infection and following TLR stimulation (56). GSK3b can be phosphorylated downstream of both ERK1/2 and p38 (57, 58). Phosphorylation of GSK3b inactivates the protein and has been shown to prevent neutrophil apoptosis following LPS stimulation (57). GSK3b inhibition has also been shown to reduce proinflammatory cytokines and increase IL-10 production by innate immune cells (56). Further, splenic macrophages from aged mice produced lower levels of inflammatory cytokines following activation of TLRs or with heat killed *S. pneumoniae* and this was associated with increased GSK3b phosphorylation downstream of PI3K and AKT activation (59).

Here we also observed increased basal expression of p38 in PMNs isolated from aged mice. This is similar to what we observed with increased basal ERK1/2 expression with age. In a dermal model of varicella zoster virus infection, older volunteers displayed reduced immune response at the site of infection when compared to young controls and this was associated with increased p38 MAPK activation following sterile injection, supporting increased p38 activation with host aging (60). Administration of oral p38 inhibitor in these aged volunteers resulted in a decrease in proinflammatory cytokines IL-6 and TNF alpha and increased the immune response to varicella zoster virus infection (60). In a dermal model of sterile inflammation resolution of inflammation was impaired in older adult volunteers but was restored to similar levels seen in young controls following administration of an oral p38 inhibitor (29). In addition to the role of p38 in inflammation with age, prior work has shown a role for p38 signaling in driving T cell senescence (61), however, inhibition of p38 in senescent CD8+ T cells increased proliferation and mitochondrial biogenesis and induced increased autophagy in these cells suggesting that p38 inhibition can reverse this senescence phenotype (62). These studies and the data presented here demonstrate that overactivation of several MAPK components drives aberrant immune responses during aging and that MAPK components can be targeted to enhance immune cell function in an aged host.

In summary, this study identified the MAPK, ERK1/2 signaling pathway to be required for pneumococcal killing by PMNs in vaccinated hosts both in mouse models and in human participants administered the pneumococcal conjugate vaccine. However, dysregulation of this pathway with host aging, including increased basal phosphorylation and reduced activation in response to antibody opsonized bacteria contributes to the age-related decline in PMN function. This work has identified a pathway that may be targeted to improve vaccine efficacy in aged hosts and is clinically relevant to human immune responses.

## ACKNWELEDGEMENTS AND FUNDING

We would like to thank Dr. Manmeet Bhalla for his critical feedback on this manuscript. This work was supported by National Institute of Health grants R01 AG068568-01A1 to ENBG and F31 AI169889-01A1 to SRS.

## AUTHOR CONTRIBUTIONS

SRS designed research, conducted research, analyzed data, and wrote paper. AR conducted research and analyzed data. ENBG designed research, wrote the paper, and had responsibility for final content. All authors read and approved the final manuscript.

