## Supplemental Figure 1 and 2 for "Basal activation of ERK1/2 blunts the antimicrobial activity of neutrophils from aged hosts against antibody opsonized *Streptococcus pneumoniae*"

### Supplemental Data

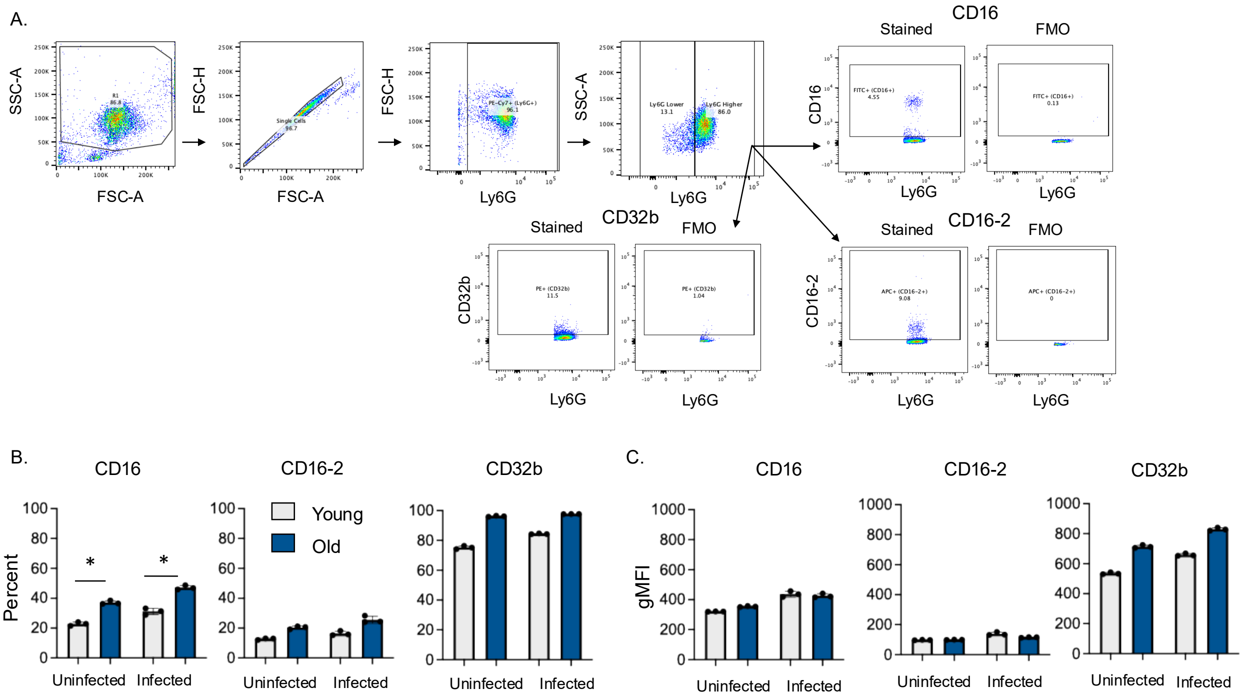


**Supplemental Figure 1: FcγR expression gating strategy and representative data.** Surface expression of FcγRs CD16, CD16-2 and CD32b on BM PMNs from young and old C57BL/6 mice at uninfected baseline and following infection with *S. pneumoniae* opsonized with immune sera. (A) Representative gating strategy. (B and C) Representative data of 1 experiment, percentage of cells expressing receptor (B) and GMFI (C) were determined by flow cytometry. * indicates *p<0.05* as determined by student's unpaired t test.

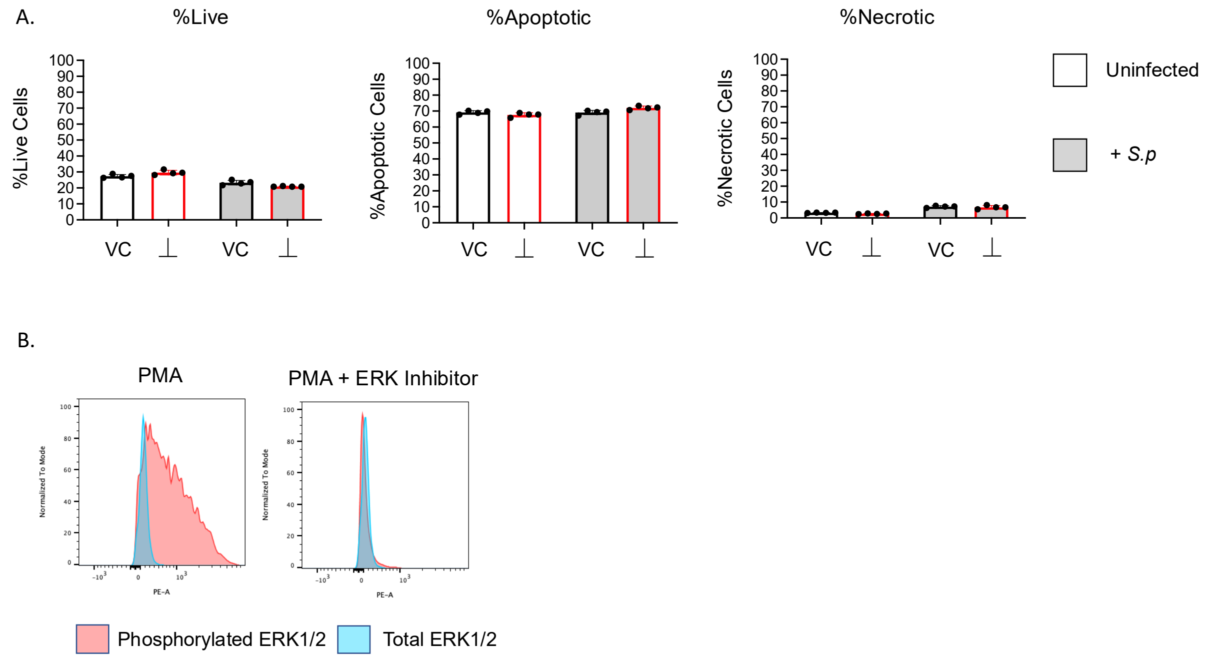


**Supplemental Figure 2: PMN viability and ERK phosphorylation in response to treatment with PD98059.** (A) PMNs isolated from the BM of C57BL/6 young mice were pretreated with specific ERK1/2 inhibitor PD98059 or VC. Following treatment, PMNs were infected with *S. pneumoniae* pre-opsonized with naïve sera. %Live, %Apoptotic, and %Necrotic cells were determined by Annexin/PI staining. (B) PMNs (Ly6G+) were stained for either phosphorylated ERK1/2 or total ERK1/2. To verify the ERK inhibitor PD98059 PMNs isolated from the BM of C57BL/6 old mice were treated with PMA alone or PMA in the presence of ERK1/2 inhibitor and phosphorylated (pink) and total (blue) ERK1/2 were measured.
